# LsGCRPred: lncRNA-SNP regulated gene expression based breast and ovarian cancer risk prediction model

**DOI:** 10.64898/2026.07.07.736696

**Authors:** Troyee Das, Gourab Das, Byapti Ghosh, Zhumur Ghosh

**Affiliations:** Department of Biological Sciences, Bose Institute, Unified Academic Campus, Kolkata-700 091, India

**Keywords:** LSNP, female cancers, cancer associated SNP, Genetic variants, Machine learning

## Abstract

Long non-coding RNAs (lncRNAs) and single nucleotide polymorphisms (SNPs) within them play crucial role in cancer susceptibility and disease outcomes. Breast and ovarian cancers, characterized by genetic heterogeneity, present significant challenges for precise diagnosis and treatment. Despite recent advancements in personalized medicine, inclusion of lncRNA-SNP (LSNP) markers into cancer risk detection panels remains limited. In this work, we put forward LSNP regulated gene expression based breast and ovarian cancer risk prediction model LsGCRPred (LSNP-Gene Interaction Based Cancer Risk Prediction Model). Notably, our approach accounts for the tissue-specificity of lncRNAs as well the benefit for individuals with predisposing conditions. Additionally, pathway analysis revealed the involvement of the LSNP interacting genes in key cancer regulating pathways. TaqMan genotyping and qPCR were performed to confirm the presence of selected LSNPs in ovarian and breast cancer cell lines along with the significant expression of the lncRNA and associated gene transcripts. These findings highlight previously overlooked genetic variants within lncRNA loci and their regulatory impact on disease outcomes, providing insights into personalized cancer diagnosis and treatment strategies. The tool LsGCRPred can be accessed as a standalone version on GitHub.

Github Link: https://github.com/zglabDIB/LsGCRPred

## 1. Introduction

Long non-coding RNA (lncRNA) molecules, surpassing 200 nucleotides in length, are predominantly transcribed by RNA polymerase II, undergoing splicing and polyadenylation processes. (Mercer, Dinger et al. 2009). Primarily situated within the nucleus, these molecules exhibit multifaceted roles in cellular mechanisms, notably implicated in cancer progression(Wilusz, Sunwoo et al. 2009, Rinn and Chang 2012).Single Nucleotide Polymorphisms (SNPs) often found within non-coding genomic regions exert significant influence on disease susceptibility, particularly in cancer contexts(Reich, Gabriel et al. 2003, Erichsen and Chanock 2004).

SNPs may influence the expression of disease-associated lncRNAs(Kulkarni, Lied et al. 2019), impact splice sites leading to the generation of alternative splice variants with modified functionality. Additionally, they can bring alterations in secondary structures such as hairpin loops which have the potential to modify RNA folding and influence interactions with target proteins or transcripts(Aznaourova, Schmerer et al. 2020).

It is essential to note that certain clinical conditions in breast and ovary increases the risk of their respective cancers. Atypical Hyperplasia and Multiple Papillomas can increase the risk of breast cancer if left untreated or poorly managed(Ali-Fehmi, Carolin et al. 2003, Al Sarakbi, Worku et al. 2006, Hartmann, Radisky et al. 2014, Hartmann, Degnim et al. 2015). Similarly, patients with Endometriosis (a gynecological condition where tissue similar to the endometrial lining is present beyond the uterus, often found on the ovaries) or Polycystic Ovarian Syndrome (PCOS) are at increased risk of developing ovarian cancer(Carmina and Lobo 1999, Daniilidis and Dinas 2009, Brilhante, Augusto et al. 2017, Murakami, Kotani et al. 2020). These are influenced by various factors including age, reproductive history and lifestyle. Discrete works have also identified shared genetic mutation, copy number alteration, protein expression and active signalling pathways(Kader, Elder et al. 2020, Throwba, Unnikrishnan et al. 2022) which underscore the complex interplay between benign conditions and the development of cancers. In this context, relying solely on morphological changes observed in tissue biopsies from such benign lesions may be insufficient for accurate risk assessment(Hirahata, Ul Quraish et al. 2022). A more comprehensive approach that incorporates genetic and molecular profiling is essential to uncover the underlying mechanisms driving malignant transformation.

Consistent with the multifactorial nature of disease progression discussed above, breast and ovarian cancers display significant heterogeneity. Genetic alterations contribute to metabolic diversity and altered interactions among molecular partners even within tumors of the same tissue origin(Kim and DeBerardinis 2019). This heterogeneity plays a crucial role in determining therapeutic susceptibilities and has the potential to predict clinical outcomes. Recent advancements include not only germline DNA screening for BRCA1/2 mutations but also tumor-level assessment often through Next-Generation Sequencing (NGS) for personalized treatment(Santana Dos Santos, Lallemand et al. 2020). Accurate characterization of tumors and immune microenvironments through transcriptome sequencing has become essential for effective personalized cancer treatment. With the advent of tissue specific lncRNAs, lncRNA-SNP (LSNP) markers are garnering attention in recent times(Minotti, Agnoletto et al. 2018, Abdi and Latifi-Navid 2022) with individual reports depicting the role of SNPs within lncRNA loci in breast and ovarian carcinoma(Hassanzarei, Hashemi et al. 2017, Khorshidi, Taheri et al. 2017, Liu, Chen et al. 2017, Saeedi and Ghorbian 2020).

Current cancer risk prediction models are primarily based on DNA-level alterations, including mutational signatures and coding variants. For instance, HRDetect identifies BRCA1/2 deficiency using genome-wide mutational patterns(Davies, Glodzik et al. 2017). Machine learning (ML) approaches have been developed to predict the pathogenicity of missense variants(Capriotti and Altman 2011). More recent studies have expanded variant detection from RNA sequencing data, enabling the identification of somatic mutations across cancers(Tang, Liu et al. 2024). SNP-based models have also been applied to predict disease susceptibility in complex disorders(Ramazanova, Matkarimov et al. 2026). These approaches largely focus on coding regions or aggregate genomic signals, with limited consideration of regulatory non-coding elements and their functional interactions.

Despite these advances, a significant gap remains in the incorporation of LSNP markers into breast and ovarian cancer risk detection panels. This is particularly relevant for patients having aberrant breast and ovarian clinical conditions which acts as prelude in developing cancer at a later stage. This work focuses towards looking into the presence of tissue specific LSNPs along with their interaction with protein coding genes within biopsy samples from breast or ovarian tissues of such patients.

In this study, the work has been done in two phases: First, utilizing publicly available gene expression datasets of cancer patients, we have identified exclusive LSNPs in breast and ovarian cancer system. Next, we implemented ML techniques to develop breast and ovarian cancer risk prediction model named LsGCRPred by integrating tissue-specific LSNP markers from tissues of epithelial-origin. This includes clinical conditions that may serve as precursors leading towards cancer development.

This has been followed by wet lab validation of the most important LSNPs along with the expression validation of the corresponding lncRNAs and their interacting gene partners. Our findings highlight the significance of previously overlooked genetic variants within lncRNA loci and their regulatory influence on interacting partners, ultimately dictating cancer risk outcomes.

## 2. Materials and Methods

### 2.1. Raw Data Collection

Raw RNA-seq data for patients with ovarian epithelial and breast cancers (including Luminal, TNBC and Her2 subtypes), along with their respective healthy or adjacent normal counterparts, were sourced from NCBI GEO(Barrett and Edgar 2006)and NCBI SRA(Leinonen, Sugawara et al. 2011). Annotated SNP data for the Human genome (GRCh38) were retrieved from NCBI dbSNP (version 146)(Sherry, Ward et al. 2001). RNA-seq data corresponding to 345 breast carcinoma, 32 normal control and 101 adjacent-normal control samples were taken. For ovarian cancer, a total of RNA-seq data corresponding to 299 ovarian epithelial carcinoma, along with 51 non-cancerous control samples of ovary and fallopian tube tissue of epithelial origin, were obtained. **Table 1** shows the subtype specific distribution of cancer and non-cancerous tissue samples.

**Table 1:**
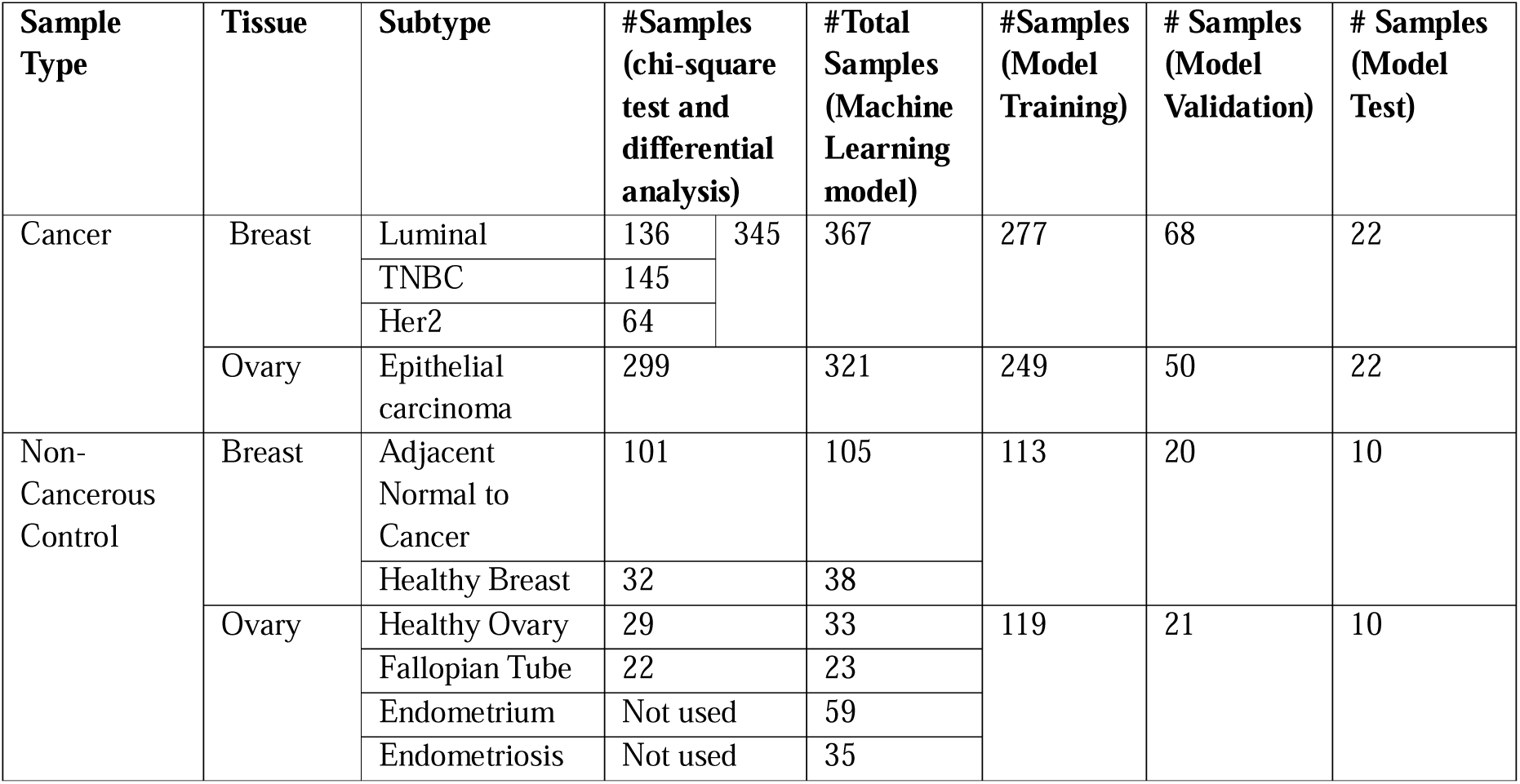
Input dataset corresponding to subtype specific breast, ovarian cancer tissues and their control counterparts respectively. This entire data has been grouped into: *Training*, *Validation* and *Test* datasets.

### 2.2. Data pre-processing and SNP detection

FastQCv.0.11.7 was employed for assessment of read quality for each sample, retaining only those with a quality score exceeding 30. Adapter sequences were then trimmed using Cutadapt v1.16(Martin 2011). Subsequently, the paired-end raw sequence reads were aligned to the Human reference genome (hg38) through HISAT22.1.0 (Pertea, Kim et al. 2016). The resulting BAM files underwent organization and indexing with SAMtools 0.1.19(Li, Handsaker et al. 2009). Before initiating variant calling, the BAM files underwent preprocessing using Opossum 0.2(Oikkonen and Lise 2017), a tool tailored for quality control procedures. This preprocessing step included the removal of duplicate or poorly mapped reads and secondary alignments, as well as the merging of overlapping reads. Opossum seamlessly integrates with the Variant Caller Platypus(Rimmer, Phan et al. 2014), which was then employed for SNP and indel detection with default parameters. Platypus is renowned for its speed and sensitivity, placing it at par with other leading variant callers in the field. Variants with a QUALITY status of “PASSED” and more than 5 reads retaining the variants (Platypus filter ‘TR’ or #Total number of reads containing this variant) were selected from the output. Following the variant filtering process, annotation of the variants were executed using dbSNP (version 151) through the “bcftools(v1.9) annotate” tool (Li, Handsaker et al. 2009).

### 2.3. Screening exclusive breast and ovarian cancer LSNPs

An SNP matrix file was generated incorporating data of 345 breast cancer patients and 299 ovarian cancer patients to identify exclusive SNPs in each of these two cancers. Thechi-square test was performed with a significance threshold value of p≤ 5e-12. The selected SNPs underwent further filtering based on the following criterions: (a) Mapping SNPs within lncRNA loci not overlapping with any protein coding region was done using bedtools(v2.26.0) Intersect(Quinlan and Hall 2010).(b)The capability of an SNP to disrupt the lncRNA secondary structure was assessed using RNAsnp(Sabarinathan, Tafer et al. 2013), which predicts the effect of SNP on local RNA secondary structure, with a p-value threshold < 0.2 (Sabarinathan, Tafer et al. 2013) indicating significant structural change. (c) The common variants were filtered out and among them only low-frequency variants, which are more prone to be linked with disease risk (Kido, Sikora-Wohlfeld et al. 2018) were screened out.

### 2.4. Detection of differentially expressed LSNPs

Using both cancer and control BAM files, StringTie v1.3.4d (Pertea, Kim et al. 2016)was used to assemble transcripts. For lncRNAs, transcript information have been retrieved from our in-house database lncRBaseV.2(Das, Deb et al. 2021) which offers a comprehensive non-redundant list of lncRNAs from various sources. The lncRNAs encompassing the selected SNPs are checked for their expression and filtered out. Further, for protein-coding genes, annotations were retrieved from Gencode(Frankish, Diekhans et al. 2021).

Differential gene expression analysis of patient samples were performed using Ballgown(Pertea, Kim et al. 2016). In the context of breast cancer, both healthy breast biopsy tissue and non-involved tissue adjacent to the tumor were utilized as control samples. Conversely for ovarian cancer, healthy epithelial tissue originating from the ovaries and fallopian tubes was considered as the control group. The stat test function within Ballgown addressed various considerations, including potential batch effects arising from multiple data sources. Differentially regulated mRNAs were identified applying a fold change (FC) cut-off of ≥2 along with a significance threshold of p-value ≤ 0.05. The differential expression status of the selected lncRNAs were also checked with a FC cut-off of ≥1.5 and p-value ≤ 0.05.

### 2.5. Screening LSNP-interacting gene partners

Three modes of interaction of lncRNAs were considered while screening the LSNP-interacting genes which are as follows:

(a) *Expression correlation:* Cis and trans correlation analyses were conducted using cancer datasets to identify robust lncRNA target genes. For cis-pairs, the differentially expressed (DE) coding genes located within±20kb of an lncRNA loci were considered. Trans co-expression analysis between DE mRNA and lncRNA was conducted employing a screening criterion of rho (Spearman coefficient) ≥ 0.7 and p-value ≤ 0.05. This analysis was performed using the rcorr function from the Hmisc library in R, which can be accessed at https://cran.r-project.org/web/packages/Hmisc.

Additionally, in order to explore the tissue-specific functions of lncRNAs, “guilt by association” method has been adopted(Lefever, Anckaert et al. 2017, Guo, Jian et al. 2019). We accessed datasets containing 1569 breast cancer samples and 507 ovarian cancer samples from the TCGA database through the GDC portal (https://portal.gdc.cancer.gov/). Employing the Spearman method in R with a threshold of Spearman coefficient>0.5 and p≤0.01, we conducted correlation analyses between the lncRNAs and all 19,962 coding genes within the respective cancer samples. Following this, we scrutinized the differential expression status of the correlated genes in our analysis, including those which were DE in the corresponding cancers with respect to their control group.

(b) *RNA-RNA interaction information*: StarBaseV2.0(Li, Liu et al. 2014)has been considered to collect lncRNA-mRNA interacting pairs identified from high-throughput sequencing data of RNA-RNA interactome, such as LIGR-Seq(Sharma, Sterne-Weiler et al. 2016), PARIS(Lu, Gong et al. 2018), SPLASH(Chaung, Baharav et al. 2023).

(c) Competing endogenous mode of interaction: Following the ceRNA theory which posits that lncRNAs can act as endogenous “sponges” to regulate mRNA expression by sequestering miRNAs, we constructed a lncRNA–miRNA–mRNA network. Human miRNA sequences were obtained from miRBase(Griffiths-Jones, Grocock et al. 2006). Point mutations were introduced at the selected lncRNA loci corresponding to the mapped SNPs to investigate the creation of new miRNA binding sites (8mers, 7mer-m8 and 7mer-A1) using TargetScan Release 8(Grimson, Farh et al. 2007). The function of these miRNAs were explored in the literature, and DE mRNAs with potential target sites within the 3’ UTR region for these miRNAs were identified.

By consolidating the outputs from the four aforementioned approaches, a distinct set of interacting partner genes have been identified for each screened LSNPs. The expression pattern of these interacting genes have been used as the backbone to serve as the feature set for developing the prediction model for cancer risk prediction.

### 2.6. ML based analysis and developing cancer risk prediction model

The objective behind developing this prediction model was not only to distinguish cancerous from healthy tissues but also to evaluate whether the identified LSNP-associated signatures could detect epithelial tissue-derived clinical conditions that are associated with an elevated risk of subsequent cancer development. Therefore, to represent clinically relevant non-cancerous ovary-associated conditions, 59 healthy endometrial tissue (HEMT) samples and 30 endometriosis (EMT) samples were incorporated into the model. Endometriosis has been widely recognized as a precursor condition associated with an increased risk of specific ovarian cancer subtypes, making it a suitable cohort for assessing the predictive capability of the model beyond established malignancies(Brilhante, Augusto et al. 2017). PCOS samples were not considered because the condition predominantly involves granulosa/theca cell dysfunction and does not directly represent a precursor state of epithelial-origin, relevant to ovarian epithelial carcinoma. Comparable publicly available transcriptomic datasets representing benign breast epithelial lesions with sufficient sample numbers and associated clinical information were not available during model development. Thus analogous breast precursor conditions could not be incorporated into the breast cancer model. These, along with the datasets mentioned in **Table 1** were divided into *training* and *validation* sets. Additional independent datasets, not used during model development, were further incorporated as *Test* to evaluate model performance on unseen data. **Table 1** summarizes the entire set of training, validation and test data (total samples: 981), and detailed dataset information is provided in **Supplementary File 1**.

By considering the expression value of the genes associated with specific LSNPs as features (as shown to be screened from the previous steps), we have incorporated the simplest linear model in forms of logistic regression to segregate the normal samples from the cancerous one. The workflow has been executed in the following way:

(a) **Initial Model Building:** Using the specific set of genes interacting with each LSNP, we developed an initial logistic regression model. This approach generated feature importance scores in the form of positive and negative coefficients (feature weights). These scores were then used to select features in the subsequent stages.

(b) **Feature Selection by Recursive Feature Combinatorial Addition Method and final model building:** For each LSNP-associated positive feature gene, negatively weighted features were added stepwise in a cumulative manner during the feature selection. At each step, a logistic regression model was trained and evaluated on the validation dataset to assess its predictive performance. The goal of this process was to identify the subset of interacting genes that most effectively distinguished cancer from non-cancer samples. The selected feature gene set was then used to optimize the final logistic regression classification model. We have utilized the python based Scikit-learn (https://scikit-learn.org/stable/index.html) tool to create the ML model.

### 2.7. Pathway Analysis

The genes constituting the final feature set for each LSNP after model training have been subjected to Ingenuity Pathway Analysis (IPA) (www.qiagenbioinformatics.com/products/ingenuity-pathway-analysis/) to check their functional involvement in specific cancer.

### 2.8. Breast and ovarian cancer cell lines

The selected set of DE lncRNAs and their corresponding SNPs were chosen for experimental validation through wet bench procedures. Breast cancer cell lines (MCF7, T47D, and ZR751) and ovarian cancer cell lines (SKOV3, OVCAR3 and PA1) were specifically selected for this validation. T47D, ZR751 and PA1 cell lines were sourced from the National Centre for Cell Science (NCCS), Pune, while MCF7 (Cat No. 300273), OVCAR3(Cat No. 300307) and SKOV3(Cat No. 300342) were acquired from the then Cell Line Service (CLS)(now known as Cytion). OVCAR3, ZR751 and T47D were cultured in RPMI-1640 medium, MCF7 was cultured in DMEM medium, and SKOV3 was cultured in DMEM: Ham’s F12 medium. All culture media contained 10% fetal bovine serum (FBS, Invitrogen) and 1% penicillin/streptomycin (Invitrogen). The cells were incubated at 37°C under 5% CO and allowed to grow until they became confluent. **Supplementary Fig. 1** illustrates the cellular condition at confluency.

### 2.9. Genotyping by the TaqMan PCR assay

Genomic DNA from the specified cell lines were isolated using the HiPura mammalian genomic DNA preparation Kit provided by Himedia.The Ready-to-use and Custom TaqMan® SNP Genotyping Assay mix for the selected SNPs (Catalog number:4351379, Assay ID: C 3130439_10 and Catalog number: 4331349, Assay ID: ANNMCVH respectively) were purchased from Thermo Fisher Scientific. TaqMan PCR and genotyping analyses were performed on the Applied Biosystems 7500 Real-Time PCR System following the manufacturer’s instructions. The reaction mixtures were amplified with 0.5 μl of 20X primer-probe mix, 5 μl of 2X TaqMan Universal Master Mix (Applied Biosystems), 1 μl of gDNA (10 ng/μl), and 3.5 μl of ddH2O, the total volume being 10 μl. PCR cycling conditions are as follows: one cycle at 95 °C for 10 min; 40 cycles at 95 °C for 15 s and 58 °C for 1 min. The results were analyzed using the allelic discrimination assay on the Applied Biosystems 7500 Real-Time PCR System.

### 2.10. Real-time PCR analysis

Normal breast and ovarian tissue RNA were procured from Biochain (Cat # R1234086-50 and C605167). Total RNA extraction from cell lines was carried out using the RNeasy® Mini kit (Qiagen), and complementary DNA (cDNA) synthesis was performed using the Verso cDNA Synthesis Kit (Thermo Fisher Scientific) following the manufacturer’s instructions. Our aim was to check the differential expression of the DE lncRNA bearing the SNP as well as a selected list of feature genes corresponding to each LSNP. Subsequently, qPCR for the selected lncRNAs was conducted using the SYBR Green PCR Master Mix (Applied Biosystems) on the Applied Biosystems 7500 Real-Time PCR System. Each experiment was performed in triplicate. Transcript expression levels were calculated using the 2^−ΔΔCt^ method, normalized to the expression of the endogenous control (18s). Primer sequences as summarized in **Supplementary File 2** were designed using NCBI primer blast and Sigma Oligo Evaluator and were ordered from Sigma-Aldrich. The entire workflow of the study is illustrated in **Fig. 1**.

**Fig. 1:**
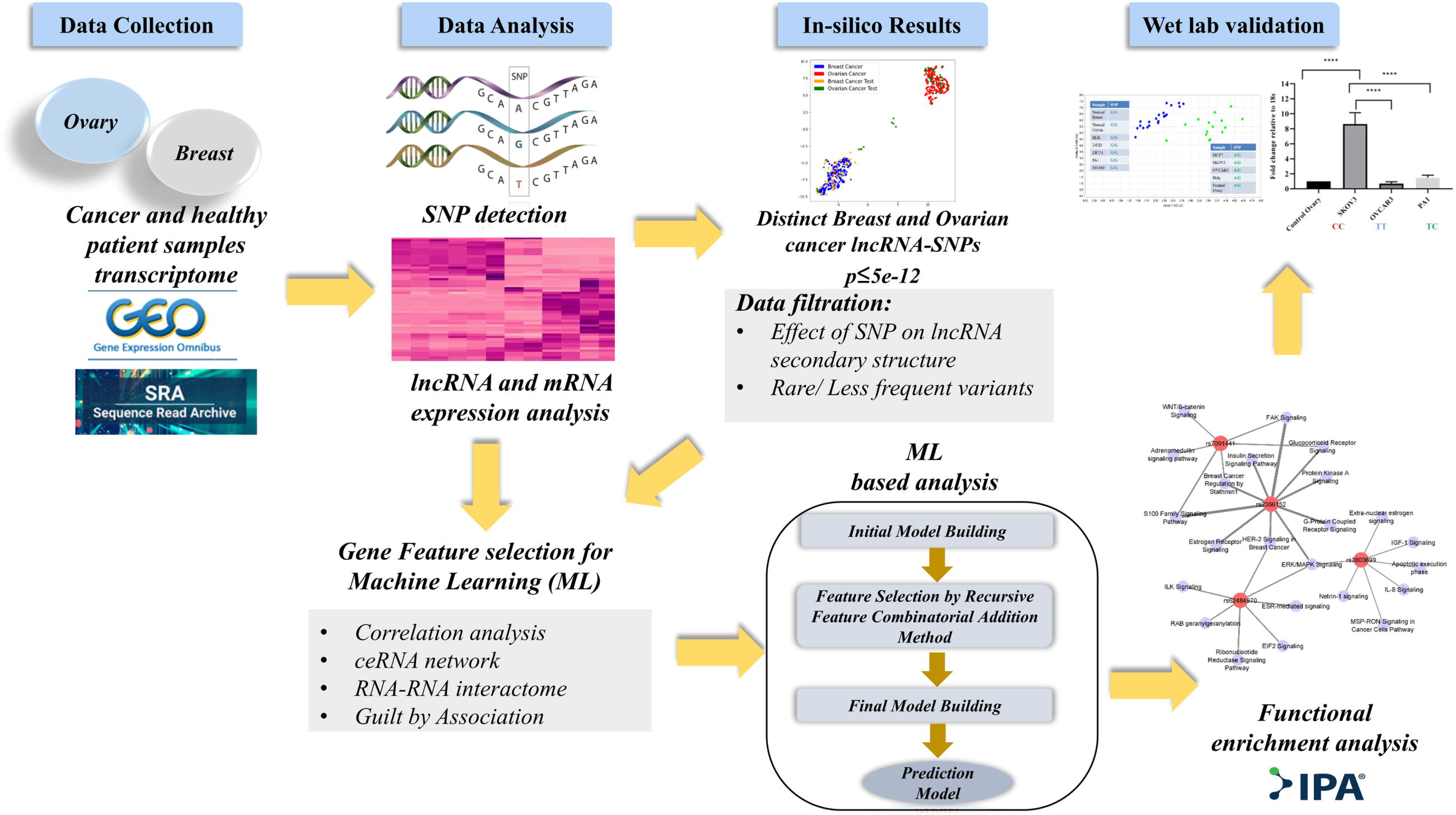
The workflow of the entire study, illustrating the sequential steps involved: starting with data collection, followed by processing and filtration, feature selection, model training, functional analysis and validation.

## 3. Results and Discussion

### 3.1. Selection of low frequency variants exclusive for breast and ovarian cancer patients

After variant calling and filtration across patient transcriptomes, a chi-square test with a stringent significance threshold of p≤ 5e-12 identified a total of 5777 distinct SNPs predominant in either breast or ovarian cancer patients. Subsequent mapping of these SNPs to lncRNA loci that do not overlap with any protein-coding regions identified a subset of lncRNA-associated variants. We then selected only those variants predicted to disrupt the secondary structure of the corresponding lncRNAs reducing the total number to 270.

Exploring allele frequencies across diverse populations revealed that most SNPs were common variants prevalent in the population. This prevalence could be attributed to natural selection where certain SNP alleles confer resistance to diseases or offer survival advantages. Additionally, random fluctuations in allele frequencies over time may contribute to their increased prevalence, despite lacking selective advantages.

While common variants are implicated in disease susceptibility, our focus shifted towards *rare* or *low-frequency variants* in disease risk association studies. These variants hold significant interest due to their potential to exert a larger effect on disease risk compared to common variants. They exhibit population specificity and are less likely to be in linkage disequilibrium with nearby markers thereby directly tagging specific functional variants or genes. Consequently, we excluded common variants from our study allowing us to concentrate on less-frequent variants potentially pivotal in elucidating the genetic factors underlying disease susceptibility.

We finally got 16 less-frequent LSNPs. We categorized them as breast and/or ovarian cancer specific, based on their frequency of occurrence within the patient samples of specific cancer type (detailed data is provided in **Supplementary File 3**). Finally, we checked the differential status of the associated lncRNAs in their respective cancer systems with respect to normal counterparts, ultimately identifying a final set of 9 LSNPs with FC cut-off of ≥1.5 and p-value ≤ 0.05 as mentioned in Section 2.4.

### 3.2. LSNP mediated effects on gene regulation for improved model efficiency

As mentioned in section 2.5 of Methodology, we investigated the presence of DE coding genes within the vicinity of the lncRNAs harboring selected LSNPs. We also identified regulatory players through guilt by association studies. In this context, guilt by association refers to inferring potential functional relationships between genes based on their co-expression patterns and shared regulatory behaviour across samples(Gillis and Pavlidis 2011). Recognizing ceRNA as one of the well-established mechanisms of lncRNA regulation of coding genes, we also carried out analysis to incorporate this aspect into our study. Additionally, physically interacting lncRNA-mRNA pairs from RNA-RNA interactome data were integrated, considering the DE status of the mRNAs in patient datasets. This approach allowed us to incorporate both the SNP itself and its effect mediated through the function of the associated lncRNAs into the prediction model. The list of LSNP interacting genes has been provided in github.

We attempted to evaluate our DE LSNPs via traditional Odds Ratios (ORs), but the severe imbalance in case–control counts yielded wide confidence intervals and nonsignificant p-values despite OR > 1 across variants. To overcome the resulting lack of statistical significance we incorporated supervised-learning based prediction model. By integrating resampling strategies (oversampling controls or under-sampling cases) and setting class_weight=“balanced”, our classifiers automatically adjusted for sample-specific skewness. Performance was then assessed using precision, recall, F1-score, and ROC AUC (**Table 2**) rather than p-values thereby capturing subtle yet significant SNPs in imbalanced data.

**Table 2:** Performance metric of LsGCRPred for breast and ovarian cancer. ROC-AUC, Accuracy and F1-Score have been provided for both Validation and Test datasets. For breast cancer: a total of 2 LSNP pairs corresponding to 2 unique SNPs and 2 unique LncRNAs, for ovarian cancer: a total of 7 LSNP pairs corresponding to 7 unique SNPs and 3 unique LncRNAs. The total number of unique genes used as model features have been shown

| SNP_ID | lncRNA | Cancer | #Feature gene set | Validation Data |  |  | Test Data |  |  |
| --- | --- | --- | --- | --- | --- | --- | --- | --- | --- |
|  |  |  |  | Accuracy | F1-Score | ROC-AUC | Accuracy | F1-Score | ROC-AUC |
| rs2366152 | HOTAIR | Breast | 175 | 89.66 | 93.13 | 95.51 | 87.5 | 91.67 | 82.73 |
| rs7091441 | LINC00993 |  | 59 | 88.51 | 92.06 | 96.52 | 87.5 | 91.67 | 95 |
| rs932501 | AL079338.1 | Ovary | 61 | 95.77 | 96.91 | 99.81 | 84.38 | 87.18 | 96.36 |
| rs2072588 | HAGLROS |  | 85 | 97.18 | 97.96 | 99.71 | 81.25 | 84.21 | 89.09 |
| rs7318592 | LINC00621 |  | 134 | 97.18 | 98 | 99.05 | 90.63 | 92.68 | 99.09 |
| rs9506960 | LINC00621 |  | 104 | 95.77 | 96.91 | 98.86 | 87.5 | 90 | 98.64 |
| rs9510420 | LINC00621 |  | 75 | 91.55 | 93.75 | 97.14 | 87.5 | 90 | 94.09 |
| rs12583808 | LINC00621 |  | 87 | 98.59 | 98.99 | 99.81 | 81.25 | 84.21 | 92.73 |
| rs60135126 | LINC00621 |  | 56 | 95.77 | 96.91 | 99.43 | 87.5 | 90 | 91.82 |

Based on this, we developed breast and ovarian cancer risk prediction models integrated within LsGCRPred, utilizing gene expression data of LSNP interacting genes. These served as feature gene set to construct the cancer risk prediction model for the selected LSNPs.

In addition to control datasets, we incorporated endometrial samples with and without endometriosis as non-cancerous controls (based on available data). Absence of publicly available transcriptomic datasets representing benign breast epithelial lesions with sufficient sample numbers compelled us to consider control and adjacent-normal tissue datasets for breast cancer analysis. While adjacent normal tissue may harbor the same SNPs as that by cancerous tissue, differences in gene expression patterns of the interacting partners can arise due to tumor-induced changes, epigenetic modifications, the tumor microenvironment, clonal selection and post-transcriptional regulation. We believe that inclusion of these aspects will enhance the robustness of our prediction model and its applicability in real-world clinical scenarios.

Following the identification of LSNP interacting genes as feature set, we generated four gene expression matrices to serve as input for each LSNP. These matrices were structured to capture distinct sample groups: (a) control/adjacent-normal/ non-cancerous samples containing the SNP of interest, (b) control/ adjacent-normal/ non-cancerous samples lacking the SNP, (c) cancer samples with the SNP and (d) cancer samples without the SNP. Each matrix represents a specific combination of genetic and disease status facilitating the training and evaluation of separate predictive models for each scenario. By organizing the data in this manner, we aim to discern the unique gene expression patterns associated with the presence or absence of each LSNP in both control and cancer samples.

Performance metrics of LsGCRPred for breast and ovarian cancer have been detailed in **Table 2**.

### 3.3. Pathway and gene set enrichment of the genes constituting the feature set reveals a significant role in breast and ovarian cancer

IPA analysis of the selected genes constituting the final feature set for each of the 2 breast LSNPs and 7 ovarian LSNPs revealed their involvement with breast and ovarian cancer respectively. **Fig. 2A and 2B** represents the proportion of the genes(in percentage) [constituting the feature set corresponding to each LSNP] which are having reported role in breast and ovarian cancers respectively.

**Fig. 2:**
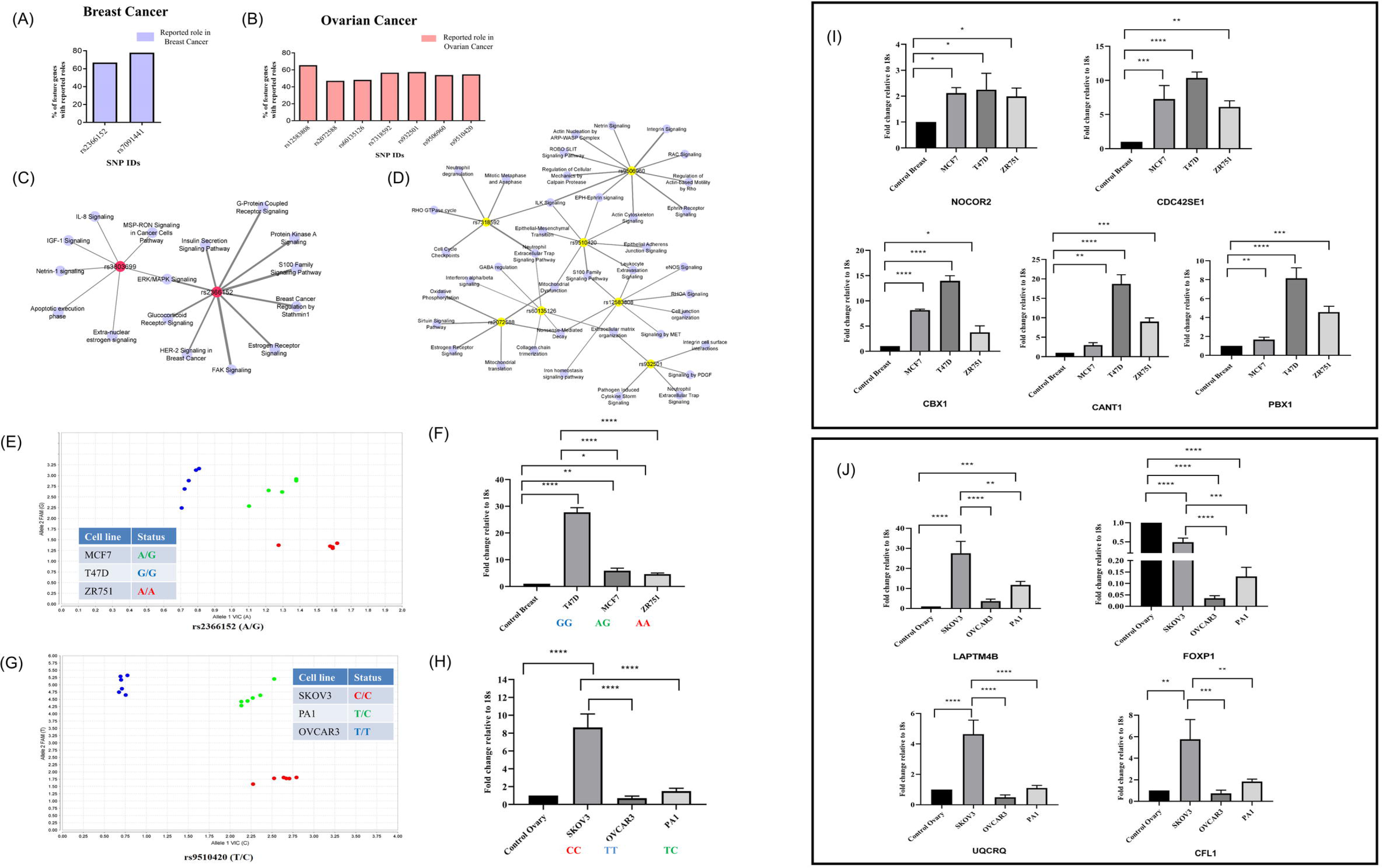
2A–2B: Bar graph illustrating the percentage of LSNP interacting genes constituting the feature set corresponding to the 2 LSNPs associated with breast cancer and 7 LSNPs associated with ovarian cancer having reported role in breast and ovarian cancer respectively; 2C–2D: Gene set enrichment analysis revealed top significant cancer related pathways corresponding to breast and ovarian cancer specific LSNP interacting genes. Breast cancer specific LSNPs are indicated in red and ovarian cancer specific LSNPs are indicated in yellow; pathways in blue and thickness of the edge reveals the no. of genes involved in each pathway 2E–2H: TaqMan genotyping assay of (E) LSNPs HOTAIR-rs2366152 and (G)LINC00621-rs9510420 and RT-qPCR expression of (F) HOTAIR and (H) LINC00621 in breast and ovarian cancer cell lines respectively For rs2366152: AA (red)-homozygous dominant allele; GG (blue)-homozygous recessive allele and AG (green) - heterozygous allele For rs9510420: TT (blue)- homozygous dominant allele; CC (red)- homozygous recessive allele and TC (green)- heterozygous allele 2I–2J: Expression validation of the target genes corresponding to HOTAIR-rs2366152 and LINC00621-rs9510420 in breast and ovarian cancer cell lines respectively.

We proceeded to analyze the top significant pathways (with p-values < 0.05) associated with each LSNP, which revealed their involvement in key cancer-regulating pathways. In the context of breast cancer, enrichment in top cancer pathways such as ERK/MAPK signalling, FAK signalling, glucocorticoid receptor signalling, HER-2 signalling in breast cancer, and the S100 Family Signalling Pathway were observed. Conversely, for ovarian cancer, pathways like estrogen receptor signalling, ILK signalling, leukocyte extravasation signalling, nonsense-mediated decay, and the S100 family signalling pathways were enriched. **Fig. 2C** and **2D** present the cytoscape network illustrating the top enriched cancer pathways involving the set of interacting genes corresponding to each LSNPs in breast and ovarian cancers respectively.

### 3.4 Performance testing of LsGCRPred on unknown test dataset

Given the absence of existing frameworks that integrate LSNPs with gene expression of the interacting genes for cancer risk prediction, direct benchmarking against established tools were not feasible. Hence, we tested the performance of LsGCRPred on clinically relevant conditions which increases the risk of these cancers. **Table 3** summarizes the prediction result (in %) in presence of ovarian cancer–associated LSNPs across two sample groups: (i) endometriosis tissue adjacent to endometrioid ovarian carcinoma and (ii) ovarian endometriosis cyst–derived carcinoma. Ideally true negative controls would consist of endometriosis samples with long-term follow-up confirming no progression to malignancy. However, to the best of our knowledge, such datasets were not available. As an alternative we utilized the data corresponding to endometriosis tissue adjacent to endometrioid ovarian carcinoma (n = 10, GSM4756719–GSM4756728). These lesions being located near the tumor yet remaining histologically non-malignant, resembles endometriotic tissue that has not undergone malignant transformation. Samples with predicted probabilities(%) ≥ 50 were classified as high-risk, whereas probabilities(%) < 50 were classified as low-risk. Our model revealed the probabilities of elevated cancer risk in terms of percentage for these samples. Interestingly, the samples (i.e. n=10, GSM4756719–GSM4756728) corresponding to endometriosis tissue adjacent to endometrioid ovarian carcinoma consistently lacked the LSNPs of interest. In contrast, analysis of ovarian endometriotic cyst–derived carcinoma samples (n =10, GSM7086123-GSM7086132) revealed the presence of multiple SNPs with positive prediction scores in most of the cases, supporting the model’s ability to distinguish between lower-risk and higher-risk disease-associated states.

**Table 3:**
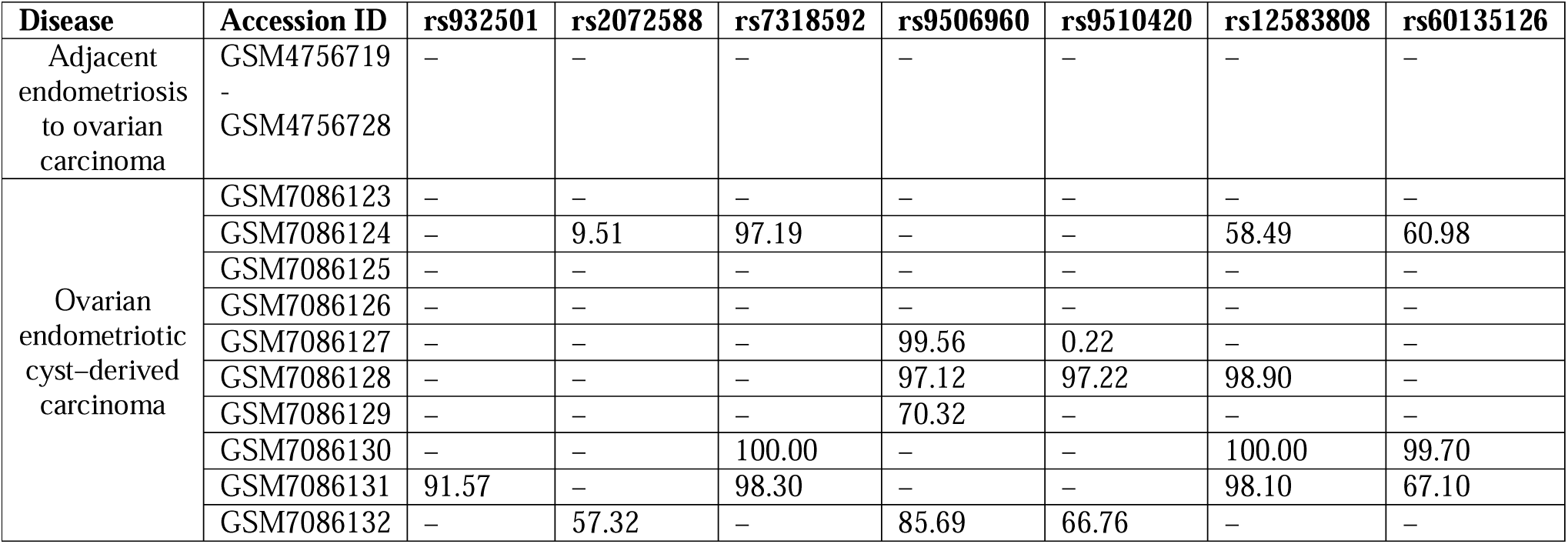
Prediction result (in percentage) across clinically relevant samples in presence of ovarian cancer specific LSNPs. ‘-’ indicates absence of the SNPs or no detectable signal. Numeric values represent model-predicted cancer risk probabilities (%) for samples in which the LSNPs were detected. The table summarizes the prediction result (in %) in presence of ovarian cancer–associated LSNPs across two sample groups: (A) endometriosis tissue adjacent to endometrioid ovarian carcinoma and (B) ovarian endometriosis cyst–derived carcinoma

Together, these results follow the expected biological trend and depicts the performance of the prediction model (for each of the LSNPs corresponding to ovarian cancer) towards assessing cancer risk in predisposing clinical conditions. A comparable evaluation was not feasible for breast cancer owing to the lack of publicly available transcriptomic datasets from clinically relevant pre-malignant or high-risk breast lesions. Future studies incorporating such datasets will enable a more comprehensive assessment of LsGCRPred in breast cancer risk stratification.

### 3.5 Screening the most promising LSNPs for wet lab validation

Reviewing our final selection of ovarian LSNPs, we made an intriguing observation: LINC00621 harbors five out of the twelve chosen SNPs. This clustering of SNPs within the locus of LINC00621 sparked our curiosity. Upon delving deeper, our literature survey unveiled LINC00621 as an onco-lncRNA in lung cancer(Wei, Yu et al. 2023). However, its involvement in ovarian cancer has not been explored. Utilizing our previous findings, where we analyzed RNA-seq data from ovarian cancer cell lines for variant detection, we investigated the status of LINC00621 SNPs in the ovarian cancer cell lines at our disposal. From the same cell line datasets, used during our previous work on ClinicL-SNP(Das, Deb et al. 2021), we detected rs9510420 in all SKOV3 cell lines and in PA1 cell line dataset, but notably absent in OVCAR3. Consequently, we selected LINC00621-rs9510420 for wet lab validation.

For breast cancer, we selected HOTAIR, a well-recognized lncRNA implicated in dysregulated functions within breast cancer(Cantile, Di Bonito et al. 2020). rs2366152, residing within HOTAIR loci, has previously been associated with colorectal cancer susceptibility in the Iranian population(Eivazi, Mirfakhraie et al. 2023) and cervical cancer susceptibility in the Polish population(Łaźniak, Sowińska et al. 2023). However, its involvement in breast cancer remains unexplored in existing literature. Through variant analysis of breast cancer cell line transcriptome data, we detected the presence of this LSNP in cell lines MCF7 and T47D but was absent in ZR751. Hence, we proceeded to validate the expression of HOTAIR-rs2366152 and its interacting genes.

### 3.6 Validating the presence of the breast cancer specific LSNPs and expression validation of the corresponding lncRNA

Taqman genotyping assay was carried out to check the presence of the LSNPs and q-PCR experiments were done to check the abundance of the lncRNA harboring the SNP. 18s was used as the endogenous control.

(a) *LSNP rs2366152 harbored by the lncRNA HOTAIR*: In line with our in-silico analysis, results of TaqMan genotyping assay (as illustrated in **Fig. 2E)**, indicate the presence of homozygous dominant allele (AA) of rs2366152 in breast cancer cell line ZR75, homozygous recessive (GG) allele in T47D and heterozygous variants (AG) in MCF7.

We further carried out the q-PCR experiments to check the expression of lncRNA HOTAIR in 3 breast cancer cell lines(viz. T47D, MCF&ZR751) with their corresponding normal counterpart as control samples, as illustrated in **Fig. 2F**. Comparison of expression profiles among cell lines carrying homozygous and heterozygous genotypes was performed to assess potential allele-dosage effects of the variant. Functional variants often show genotype-dependent molecular phenotypes, with different effects observed between homozygous and heterozygous carriers(Albert and Kruglyak 2015). q-PCR analysis revealed a statistically significant upregulation of the lncRNA in all the three cell lines with respect to that in control samples. However, cell line T47D, which carries the homozygous allele (GG), shows significantly higher upregulation compared to the others.

(b) *LSNP rs9510420 harbored by the lncRNA LINC00621*: The result from the TaqMan genotyping assay, have been illustrated in **Fig. 2G**. It indicates the presence of homozygous dominant allele (TT) corresponding to rs9510420 in control ovary and PA1 line, homozygous recessive (CC) allele in SKOV3 and heterozygous variants (CT) in PA1.

The differential expression analysis of the lncRNA LINC00621 in the 3 ovarian cancer cell lines (viz. SKOV3, OVCAR3 and PA1) with their corresponding normal counterpart, as illustrated in **Fig. 2H** revealed a statistically significant increase in the expression of the lncRNA in the SKOV3 cell line harboring the SNP (CC). Additionally, we observed a marginal increase in its expression in PA1 carrying the heterozygous allele (TC) and a slight decrease in OVCAR3 carrying the homozygous dominant allele (TT), though these findings did not reach statistical significance in our analysis.

Here, we note an increase in the expression levels of LINC00621 in the cell line harboring SNP rs10425267, implies the likelihood of this genetic variant impacting the expression pattern of the lncRNA within the ovarian cancer system.

### 3.7 Expression validation of the interacting genes corresponding to the selected LSNPs

We next proceeded to validate a selected set of interacting genes from the feature gene set of rs2366152-HOTAIR (**Fig. 2I**) and rs9510420-LINC00621(**Fig. 2J)** based on their expression being influenced by the presence of LSNPs as predicted by LsGCRPred as well as their involvement in pathways that could drive the benign condition towards carcinoma(**Fig. 3A–B** and functional details in **Supplementary File 4**). In breast cancer, we observed upregulation of five genes (CBX1, PBX1, CANT1, NCOR2 and CDC42SE1) across all breast cancer cell lines, with significantly higher expression in T47D cells harboring the homozygous recessive variant (**Fig. 2I**). Additionally, RNA-RNA interactome data(Li, Liu et al. 2014) showed physical interactions of three of these genes (CANT1, NCOR2 and CDC42SE1) with HOTAIR.

**Fig. 3:**
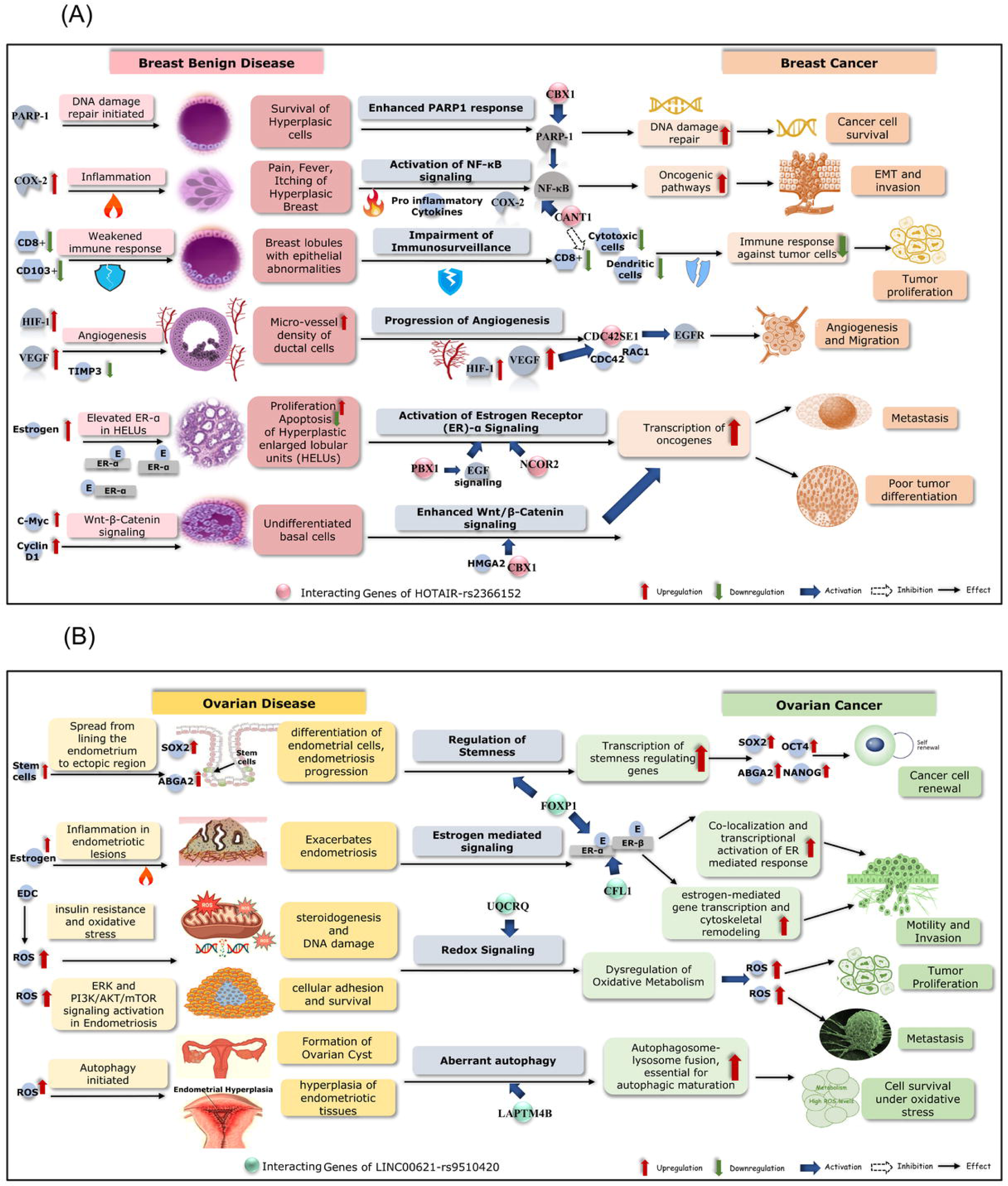
Role of the genes interacting with (A) HOTAIR-rs2366152 that drives the progression of benign breast disease to breast carcinoma and (B) LINC00621-rs9510420 that drives the progression of ovarian disease to ovarian carcinoma Abbreviation: CANT1: Calcium-activated nucleotidase-1, CBX1: Chromobox 1, CDC42: Cell division control protein 42, CDC42SE1: CDC42 small effector 1, C-Myc: MYC Proto-Oncogene, COX-2: Cyclooxygenase-2, EGF: Epidermal Growth Factor, EGFR: Epidermal Growth Factor Receptor, EMT: Epithelial-Mesenchymal Transition, ER: Estrogen Receptor, HIF-1: Hypoxia-Inducible Factor-1, HMGA2: High Mobility Group AT-Hook 2, NCOR2: Nuclear Receptor Corepressor 2, NF-κB: Nuclear Factor-kappa B, PARP1: Poly [ADP-ribose] polymerase 1, PBX1: PBX Homeobox 1, RAC1: Ras-related C3 botulinum toxin substrate 1, TIMP3: Metalloproteinase inhibitor 3, VEGF: Vascular Endothelial Growth Factor ABGA2: ATP Binding Cassette Subfamily G Member 2, CFL1: Cofilin 1, EDC: Endocrine Disrupting Chemicals, EMT: Epithelial-Mesenchymal Transition, ER: Estrogen Receptor, FOXP1: Forkhead box protein P1, LAPTM4B: Lysosomal Protein Transmembrane 4 Beta, OCT4: Octamer-binding transcription factor 3/4, ROS: Reactive Oxygen Species, SOX2: SRY-box 2, UQCRQ: Ubiquinol-Cytochrome C Reductase Complex III Subunit VII

For ovarian cancer, q-PCR analysis confirmed significantly higher expression of the four genes(FOXP1, CFL1, UQCRQ, and LAPTM4B) in SKOV3 cell line carrying homozygous recessive variant of SNP rs9510420(CC) compared to OVCAR3 (TT) and PA1 (TC) (**Fig. 2J**). Notably, RNA-RNA transcriptome data also indicated a physical interaction between FOXP1 and LINC00621 as well as CFL1 and LINC00621.

Atypical hyperplasia, a benign condition of abnormal cell proliferation in breast tissue, raises breast cancer risk three to five times and may lead to cancer in about 30% of cases within 25 years(Noorbakhsh, Koenig et al. 2022). Multiple papillomas, defined as five or more distinct papillomas in a breast tissue segment, also elevate breast cancer risk and may correlate with atypia and a higher probability of malignancy(Rella, Romanucci et al. 2022). Deep infiltrating endometriosis and ovarian endometriomas (often referred to as “chocolate cysts” due to their appearance) increase ovarian cancer risk by about 9.7 times compared to women without endometriosis(Barnard, Farland et al. 2024).The breast cancer related LSNP interacting upregulated genes are associated with NF-κBsignaling, Immunosurveillance, estrogen receptor-α signaling, wnt/β-Catenin signalling and poly ADP ribose polymerase (PARP) response(Blackmore, Karmakar et al. 2014, Lee, Liu et al. 2015, Gao, Hu et al. 2022). This is well depicted in **Fig. 3A** which shows the role of interacting genes corresponding to HOTAIR-rs2366152 that drives benign breast disease to breast carcinoma.

The validated ovarian cancer related LSNP interacting upregulated genes are associated with pathways such as estrogen signaling, angiogenesis, oxidative stress, stem cell renewal and autophagy(Li, Iglehart et al. 2012, Choi, Seo et al. 2016, Kari, Kandhavelu et al. 2023). This is well revealed in **Fig. 3B** which shows the role of the interacting genes corresponding toLINC00621-rs9510420 that drives benign ovarian diseases of epithelial origin to ovarian carcinoma. Further functional details of both these set of genes are provided in **Supplementary File 4**. These pathways are implicated in both non-cancerous and cancerous conditions of the respective tissue and are speculated to drive the transition from a benign state to malignancy. Screening patients with corresponding LSNPs and subsequently checking the expression levels of these genes could provide clues for taking early precautionary measures against breast and ovarian cancer.

## 4. Conclusion

To conclude, our study highlights the potential relationship between LSNPs, their interacting gene partners, and breast and ovarian cancer biology. Through the analysis of publicly available transcriptomic datasets and the application of ML approaches, we identified tissue-specific LSNPs and associated target genes which may serve as candidate molecular markers for cancer risk prediction. By focusing on less-frequent variants within lncRNA loci we aimed to elucidate their regulatory roles and their impact on disease susceptibility. Addressing the challenge of imbalanced case-control data, we employed ML algorithms to enhance the accuracy of SNP significance analysis. By integrating LSNP-associated gene expression signatures into the LsGCRPred framework we explored the feasibility of using these molecular features to distinguish cancerous and non-cancerous conditions across independent datasets. Pathway analysis further suggested the involvement of several cancer-associated biological processes, providing insights into potential mechanisms through which LSNP-containing lncRNAs may contribute to disease development. In addition, preliminary wet-lab validation supported the biological relevance of selected candidate markers. LsGCRPred generates probability (%)-based risk estimates (cut-off ≥50) derived from LSNP-associated molecular signatures. While additional validation is required, such risk estimates may be particularly valuable in the context of predisposing conditions, where quantitative assessment of cancer risk could aid in identifying individuals who may benefit from closer monitoring and follow-up.

Nevertheless, a major limitation of this study was the lack of publicly available relevant datasets representing benign breast epithelial conditions that might mimic precursor state for breast cancer development. Consequently, while precursor conditions for ovarian cancer such as endometriosis could be incorporated into the model, a similar evaluation could not be performed for breast cancer due to lack of available data. In future, we plan to expand the study by incorporating larger and more diverse cohorts with particular emphasis on benign and pre-malignant breast epithelial conditions. Furthermore, given the known population-specific variation in genetic architecture and cancer susceptibility, future studies will focus on developing and evaluating region and population-specific models using geographically diverse datasets. Such efforts are expected to improve both the biological understanding and clinical applicability of LSNP-based cancer risk prediction.

Overall, our findings suggest that LSNP-associated molecular signatures represent promising candidates for further investigation in cancer risk assessment and biomarker discovery. In addition, this work provides a foundation for future studies aimed at understanding the contribution of lncRNA-associated genetic variation to breast and ovarian cancer susceptibility.

## Supporting information

https://bicresources.jcbose.ac.in/zhumur/biorxiv_suppl/LsGCRPred/download.php

## Acknowledgements

We acknowledge the Bioinformatics Centre project at Bose Institute, Kolkata sanctioned by the Department of Biotechnology, Government of India (Sanction no. BT/PR40174/BTIS/137/45/2022) as well as the Science and Engineering Research Board (SERB), Government of India, (Sanction no. EMR/2016/002005) and Council of Scientific and Industrial Research (CSIR), Government of India, for funding through CSIR-NET.

## Code availability

The code is available as Github Link: https://github.com/zglabDIB/LsGCRPred

## Supplementary Files

**Supplementary File 1:** Detailed information of the input datasets corresponding to Breast and Ovarian Cancers and their normal counterparts.

**Supplementary File 2:** Primer sequences corresponding to lncRNA and genes used in this study

**Supplementary File 3**: Final set of breast and ovarian cancer LSNP details

**Supplementary File 4**: Function of differentially regulated breast and ovarian cancer genes and associated pathways observed in benign and malignant conditions

**Supplementary Fig. 1:** Breast cancer cell lines MCF7, ZR751 and T47D; ovarian cancer cell lines SKOV3, OVCAR3 and PA1

