## Supplementary material for "LsGCRPred: lncRNA-SNP regulated gene expression based breast and ovarian cancer risk prediction model": https://bicresources.jcbose.ac.in/zhumur/biorxiv_suppl/LsGCRPred/download.php: Supplementary File3.docx

**Table 1:** Association and role of Differentially regulated Ovarian cancer genes in pathways observed in benign and malignant Ovarian conditions. Red = Up, Green = Down

| **Genes dysregulated in Cancer** | **Associated Pathways** | **Role in Non-cancerous Ovarian Disease** | **Role in Cancer** |
| --- | --- | --- | --- |
| FOXP1 | Stem cell production | **Stem cell theory** posits that the cells responsible for the regeneration of the endometrial lining during one's menstrual cycle play a role in the development of endometriosis. The spreading of these stem cells to ectopic regions can then lead to the differentiation of endometrial cells and cause endometriosis[1]. Enhanced expression of stemness related genes like SOX2 and ABGA2 have also been linked to endometriosis pathology[2, 3]. | As a transcription factor, FOXP1 regulates several key target genes that are involved in **stemness and cancer stem cell characteristics in ovarian cancer cells.** Enhanced expression of genes such as OCT4, SOX2, NANOG and ABGA2 is promoted by FOXP1, which plays a critical role in sustaining the stem-like properties of ovarian cancer cells, thereby supporting tumor progression[4]. |
|  | Estrogen signaling | Endometriotic lesions often exhibit high local concentrations of estrogen, which is primarily produced by the lesions themselves due to the upregulation of enzymes responsible for estrogen biosynthesis, such as aromatase. This local production of estrogen stimulates further growth and sustains the inflammatory environment within the lesions, thus exacerbating the condition[5, 6]. | Interactions between FOXP1 and Estrogen Receptor (ER) plays key role in the progression of ovarian cancer. Expression of both FOXP1 and ERβ decrease at mRNA level while protein expressions higher in cytoplasm. Colocalization of FOXP1, ERα, and ERβ was present in the cytoplasm, with ERβ specific co-localization with FOXP1 in the perinuclear area[7]. |
| CFL-1 |  |  | CFL1 regulates actin dynamics, promoting cancer cell motility and invasiveness, and interacts with estrogen receptors, influencing estrogen-mediated gene transcription and cytoskeletal remodeling. This role is critical for understanding how hormone-responsive cells, such as those in ovarian cancer, become more aggressive under estrogen influence[8-10]. |
| UQCRQ | Reactive Oxygen species (ROS) production | Reactive oxygen species (ROS) are recognized as critical pro-inflammatory mediators in the context of endometriosis. Increased ROS levels can activate signaling pathways such as ERK and PI3K/AKT/mTOR, facilitating the growth of endometriotic lesions through enhanced cellular adhesion, angiogenesis, and cell survival[11, 12]. | UQCRQ has been associated with oxidative metabolism, which is critical for various cancers, including ovarian cancer. Its activity directly influences redox signaling pathways by modulating the production of ROS during electron transport processes. Dysregulation of UQCRQ can result in altered redox status, contributing to oxidative stress within the cellular environment[13]. |
| LAPTM4B | Autophagy | Oxidative stress or **increase in ROS** in endometriosis can induce Aberrant **autophagy** in the eutopic endometrium and ectopic endometriotic foci, promoting the hyperplasia of endometriotic tissues and stromal cells, inhibiting apoptosis, and triggering abnormal immune responses[11, 14]. | LAPTM4B plays a crucial role in regulating autophagy in cancer cells, particularly in promoting the later stages of autophagic maturation. It facilitates the fusion of autophagosomes with lysosomes, a critical step necessary for the efficient degradation and recycling of cellular components, which is essential for cell survival under metabolic stress[15, 16]. |

**Table 2:** Association and role of Differentially regulated Breast cancer genes in pathways observed in benign and malignant Breast conditions

| **Genes dysregulated in Cancer** | **Associated Pathways** | **Role in benign Breast Disease** | **Role in Cancer** |
| --- | --- | --- | --- |
| CANT1 | NF-κB signaling | Some people with hyperplasia experience fever and breast itching. Prostaglandins, which induce heat, pain, and inflammation, are produced by COX-2 (cyclooxygenase-2). Through controlling COX-2 genes, NF-κB influences proinflammatory signalling pathways in hyperplasia patients[17]. | In lung cancer, CANT1 promotes progression via the NF-κB signaling pathway. This pathway is crucial in regulating immune response, inflammation, and cell proliferation. High CANT1 expression leads to the activation of NF-κB, driving cancer cell growth and survival[18]. |
|  | Immunosurveillance | In women with benign breast disease who later developed Breast cancer, breast lobules with greater epithelial abnormalities exhibit significant reductions in cytotoxic T cells. This suggests that impaired immunosurveillance might play a role in the earliest stages of breast cancer development[19]. | In Hepato Cellular Carcinoma (HCC), CANT1 overexpression is associated with a reduction in cytotoxic cells (which kill cancer cells), dendritic cells (which present antigens and activate T cells), and CD8+ T cells. This reduction weakens the body's immune response against tumors, allowing cancer cells to proliferate and spread[20]. |
| NCOR2 | Estrogen receptor-α signaling | Hyperplastic enlarged lobular units (HELUs) are clinically significant as they are the earliest identifiable precursors to breast cancer. HELUs precede other breast abnormalities such as microcysts, usual ductal hyperplasia, and atypical ductal hyperplasia (ADH), a well-known precursor to breast cancer. The epithelial cells in HELUs exhibit high levels of nuclear estrogen receptor-α (ER-α), increased proliferation, and decreased apoptosis indicating that elevated ER-α may play a key role in HELU development[21]. Additionally, P21-activated kinase-1 phosphorylates and activates ER-α, promoting hyperplasia in mammary epithelium[22]. | NCOR2/SMRT is an independent predictor of unfavourable results in Breast Cancer[23]. NCOR2's role in the estrogen receptor (ERα) signaling (modifying the transcriptional activity of ERα or directly influencing the ERα expression) pathway is crucial for tumor growth[24]. |
| PBX1 |  |  | A prognostic marker of Breast Cancer[25], PBX1 translates epigenetic signals to mediate estrogen-induced ERα binding. By controlling the transcriptional response of ERα to epidermal growth factor (EGF) signaling, PBX1 plays a critical role in controlling a subset of EGF-ERα genes associated with aggressive breast tumors[26]. |
| CDC42SE1 | Angiogenesis | Breast Hyperplasia specimens show a significant increase in microvessel density, indicating that angiogenesis escalates as ductal cells progress from normal to hyperplastic. The expression of HIF-1α, VEGF, and TF (a transmembrane protein) begins at the onset of hyperplasia in the mammary duct, contributing to the angiogenic switch[27, 28]. | Hypoxia-inducible factor 1 (HIF-1) promotes macrophage migration in the tumor microenvironment through the NO-HIF axis, with Cdc42 and Rac1 identified as an effector that modulates the actin cytoskeleton[29]. Acting downstream of CDC42, CDC42SE1 may contribute to actin cytoskeleton organization by triggering actin filament assembly and modifying CDC42-induced alterations in cell shape (uniprot.org/uniprotkb/Q8BHL7/entry). CDC42SE1 thus along with Cdc42 may also influence the secretion of vascular endothelial growth factor (VEGF), promoting the migratory behavior of breast cancer cells and angiogenesis[30].  . |
| CBX1 | Wnt/β-Catenin signaling | When Wnt/β-catenin signalling is specifically activated in basal mammary epithelial cells, it can impact the development of the mammary gland, causing hyperplasia consisting of undifferentiated basal cells and raised levels of Myc and Cyclin D1 transcripts. Angiogenesis may also benefit from the downregulation of the angiogenesis inhibitor Timp3 expression[31, 32]. | CBX1 interacts with the HMGA2 protein, which is crucial for activating the Wnt/β-Catenin signaling pathway in HCC cancer leading to tumor vascular invasion, poor tumor differentiation, and bigger tumor sizes [33]. |
|  | poly ADP ribose polymerase (PARP) response | When single-strand DNA is damaged, PARP mediates DNA repair. In Brca1-deficient mice, PARP inhibitors have been found to prevent the development of mammary gland hyperplasia implicating its role in hyperplasic cell survival[34]. | High expression of HP1β/CBX1 mRNA was associated with a worsened relapse-free survival in all Breast Cancer patients[34]. CBX1 is associated with PARP mediated DNA damage response in Breast cancer. Breast cancer cells with reduced HP1β are found to be more vulnerable to PARP inhibitors.  [35]. |
