## Supplementary figures and images for "LsGCRPred: lncRNA-SNP regulated gene expression based breast and ovarian cancer risk prediction model"

### Supplementary Figure 1.jpg

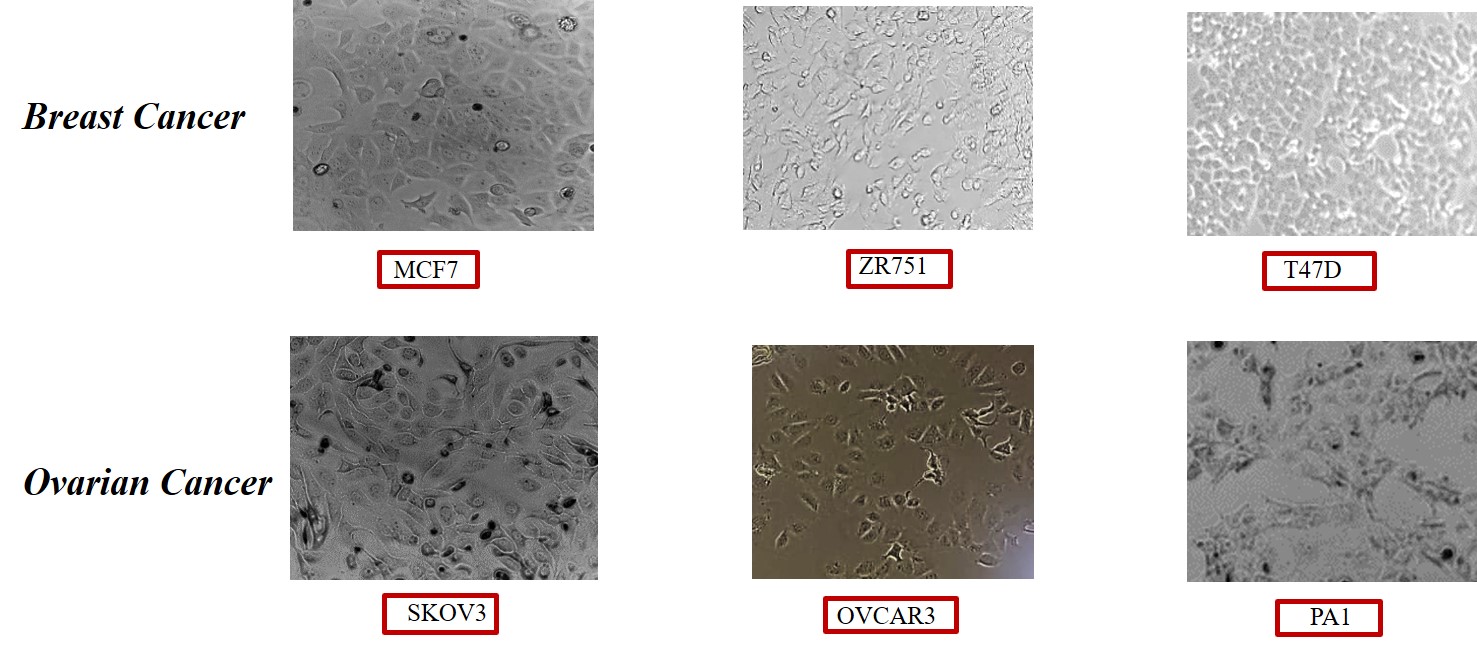
